# Decoding the Conformational Dynamics and Hyperactivity of Histone H3K36 N-Methyltransferase in Oncogenic Mutations *via* tICA and Markov State Modeling

**DOI:** 10.64898/2026.06.03.730013

**Authors:** Tejas Shah, Sahar Heidari, Jakub Rydzewski, Hedieh Torabifard

## Abstract

NSD2 is a histone methyltransferase that modifies lysine 36 in histone H3 (H3K36), playing a central role in chromatin organization and transcriptional regulation. Oncogenic mutations, such as E1099K and T1150A in NSD2, have been associated with hyperactive methylation, but the molecular mechanisms underlying this gain of function remain poorly understood. In this study, we performed all-atom molecular dynamics simulations on models of NSD2 bound to the nucleosome for the wild type (WT), E1099K, T1150A, and the E1099K/T1150A double mutant. Analysis of MD simulations reveals that the global dynamics of the enzymes remain unaltered upon mutations. The time-lagged independent component analysis (tICA) and Markov state modeling uncovered fundamental differences in free-energy landscapes among the variants. The WT NSD2 exhibited energetically and kinetically unfavorable transitions between the macrostates along with extended enzyme-substrate distances. On the other hand, the mutant systems demonstrate reduced SAM-H3K36 distances with modified energy landscapes that facilitate transitions or favor prolonged occupancy of catalytically competent states. Importantly, the mutations reorganize the network of intramolecular contacts around the catalytic site, SAM-binding pocket, and histone-binding interface, optimizing substance engagement geometry. These findings demonstrate that oncogenic mutations achieve hyperactivity through strategic reorganization of conformational dynamics rather than simple destabilization, balancing local flexibility with global stability to enhance catalytic efficiency. Our results provide mechanistic insights into NSD2 dysregulation in cancer and establish a framework to develop allosteric inhibitors that target the enzyme’s dynamic landscape.

## Introduction

The packaging of eukaryotic genomic DNA in chromatin enables the regulation of gene expression, replication, and repair while maintaining genomic integrity. The nucleosome core particle (NCP) is the fundamental unit of chromatin, which consists of approximately 147 base pairs of DNA wrapped around an octamer of histone proteins. This histone core is composed of two copies of each of histones H2A, H2B, H3, and H4. The central histone fold domains (ordered regions) form the core of the nucleosome and contribute to the architecture of the histone octamer, while the LYS and ARG-rich tails of the histone protrude from the nucleosome **(Fig. 1A)** surface and are accessible for a variety of post-translational modifications (PTMs). These PTMs form the basis of the so-called “histone-code”, which plays a vital role in the epigenetic regulation of gene expression and chromatin dynamics. ^1–10^

**Figure 1:**
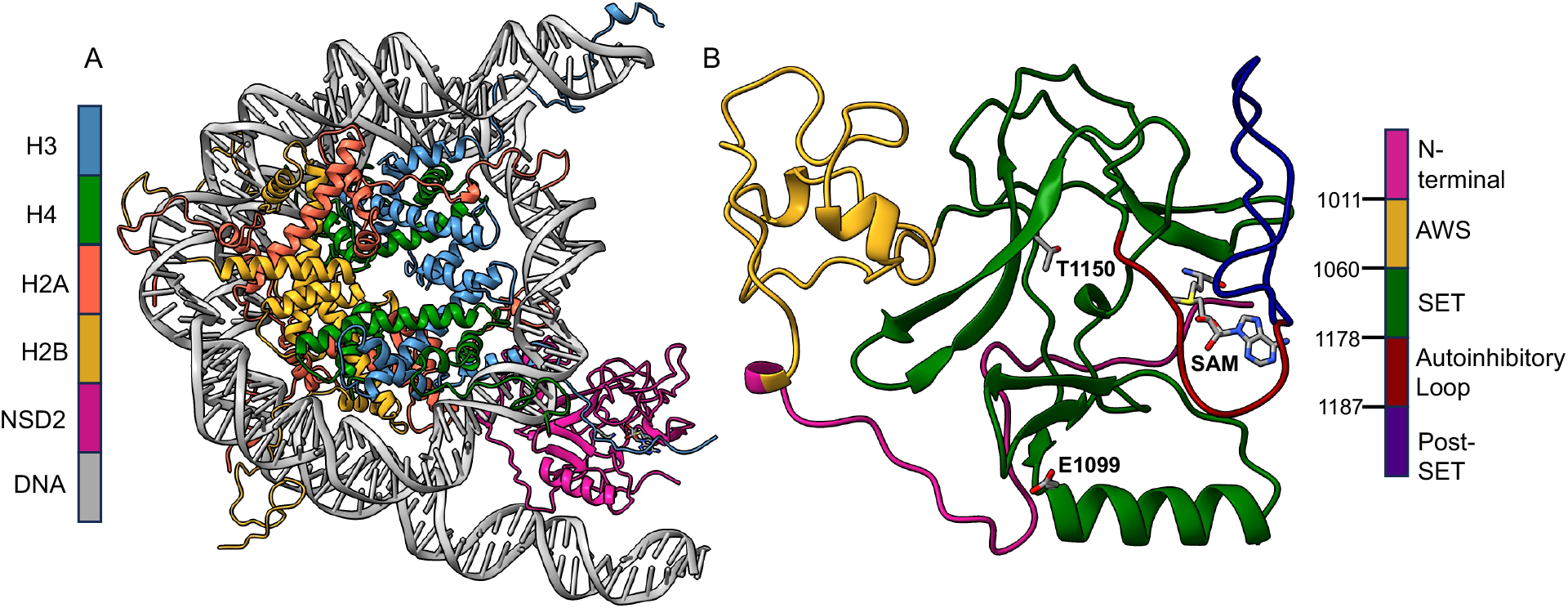
(A) Structure of nucleosome core particle with histone tails (except N-terminal of substrate H3) with NSD2. Histones H3 in blue; H4 in green; H2A in orange; H2B in yellow; and NSD2 is in magenta. (B) Structure of NSD2 with catalytic domains represented in different colors. The mutation sites E1099 and T1150, as well as SAM, are represented as licorice. Here, N-terminal is shown in magenta; AWS domain in yellow; SET domain in green; autoinhibitory loop in red; post-SET domain in violet.

Common histone PTMs include acetylation, methylation, phosphorylation, ubiquitination, and ADP-ribosylation. These modifications are regulated by specific enzymes that either add (writers), remove (erasers), or recognize (readers) the specific chemical groups. ^10–12^ Among these PTMs, methylation events, and, specifically, lysine methylation events, are remarkably versatile chemical modifications. Lysine methylation occurs in mono-, di-, and trimethylated forms, each conferring distinct chromatin states and biological functions. ^9,13,14^ For example, H3K36me1 is enriched in gene bodies and enhancers, H3K36me2 regulates DNA methylation and replication timing, while H3K36me3 is closely associated with active transcription elongation and suppression of cryptic transcription initiation.^9,10,15,16^

The role of nuclear receptor-binding SET domain protein 2 (NSD2, also known as WHSC1/MMSET) is histone H3 lysine 36 (H3K36) methyltransferase and in normal physiology has been linked to the regulation of gene expression programs during development and differentiation. ^17^ However, aberrant expression and mutations of NSD2 have been implicated in a range of pathological conditions, particularly hematologic malignancies and solid tumors. The t(4;14) chromosomal translocation, frequently observed in multiple myeloma, results in overexpression of NSD2 and leads to genome-wide enrichment of H3K36me2, which is believed to reprogram chromatin and transcriptional landscapes in favor of oncogenesis. ^1,2,9,18,19^ Furthermore, somatic point mutations (such as E1099K and/or T1150A) within the SET **(Fig. 1B)** domain of NSD2 have been identified in pediatric acute lymphoblastic leukemia and lead to hyperactivity of methyltransferase. ^13,18,20^ These findings underscore the pathological significance of NSD2 and highlight it as a potential therapeutic target.

Previous studies of SET domain methyltransferases have revealed diverse mechanisms for substrate engagement and catalytic specificity. The structures of SET7/9, SETD8, and G9a, for instance, have illustrated how these enzymes achieve specific lysine recognition and product specificity through coordinated interactions within the active site and associated regulatory domains. ^1,21–24^ The autoinhibitory loop **(Fig. 1B)** identified in SETD2, NSD1, and NSD2 highlights an evolutionarily conserved mechanism where interamolecular interactions suppress catalytic activity until substrate engagement is achieved.^1,20,25,26^

Despite its established importance, the mechanisms underlying how NSD2 achieves substrate specificity and catalyzes methylation within the complex chromatin environment remain incompletely understood. Several studies have employed molecular dynamics (MD) simulations to provide insights into key interactions, cofactor orientation, loop flexibility, and the role of oncogenic mutations in modulating methylation efficiency. However, these studies have been limited to either NSD2 alone or to NSD2 with the short H3 peptide.^9,26,27^ Nucleosome is a large, dynamic, and asymmetric substrate, and its recognition by chromatin-modifying enzymes, such as NSD2, involves multiple layers of regulation. As such, findings from the NSD2-H3 peptide complex may not capture the complete picture of the regulatory and structural constraints that govern NSD2 activity *in vivo*.

Recent cryo-EM and X-ray structures of NSD methyltransferases bound to nucleosome substrates have begun to reveal how these enzymes bind to H3 and facilitate access to the buried lysine side chains.^18,28,29^ The nucleosome-bound cryo-EM structure of NSD2 has been recently reported (PDB IDs: 7CRO^26^ and 7E8D^20^), providing the engagement and conformation of histone H3 for methyl transfer. We employed these cryo-EM structures to run MD simulations and used time-lagged independent component analysis (tICA) and Markov state modeling to understand the impact of the nucleosome, the underlying mechanisms of NSD2-mediated H3K36 methylation, and the relationship between mutant hyperactivity and the conformational landscape. The trajectories were then analyzed to quantify the conformational changes in NSD2, identify key enzyme-substrate interactions, and assess the energetic landscape of enzyme activity. This work builds on recent findings and explores the potential impact of nucleosome in dictating enzyme activity that may inform the development of selective inhibitors targeting NSD2 in anticancer therapy.

## Methods

### Model Preparation

Structures of NCP with NSD2 were reported by Sato et al.^20^ (PDB 7E8D) and Li et al.^26^ (PDB 7CRO) in 2021. Both structures contain a single copy of the NSD2 enzyme and the NCP complex, but lack the histone tails. The structure reported by Sato et al.^20^ (PDB 7E8D) contains the S-adenosyl methionine (SAM) analog, sinefungin. Initial attempts to model 7E8D by replacing sinefungin with SAM were unsuccessful; during production, SAM was found to leave the binding pocket. Hence, we opted to perform MD simulations on 7CRO.^26^ The missing residues of the histone tails were modeled by homology modeling using the SWISS model^30^ web server. All histones were separated and threaded into the template PDB (1KX5^31^) to achieve complete histone structures. The H2A tails of the histone were still missing the four residues at the N-terminal. These residues were added using the PyMol^32^ program, and the loop of the obtained histone was refined on the Modloop web server.^33^ Although this provided complete histone tails for the model, the H3 tail modeled with a 1KX5 template passes through the NSD2 enzyme and disrupts its structure upon simulation. To avoid this, the histone H3 at the active site was modeled using cryo-EM coordinates reported by Li et al.,^26^ which lacked the ≈25 residues at the N-terminal.

The protonation states of the titratable residues were obtained by H++.^34–36^ The H3 LYS36, however, was converted to its neutral form to allow methylation. The protein and nucleic acids were described using ff14sb^37^ and DNAOL15^38^ parameters available in AMBER,^39^ respectively. The improved force field parameters of SAM were obtained from Bame et al.^40,41^ The NSD2 system contains three zinc atoms; two are shared by seven CYS residues. Since the inherent Zinc Amber Force Field (ZAFF^42^) only supports tetrahedrally coordinated zinc atoms, the force field parameters for the seven Cys-two Zinc system were obtained using the `MCPB.py` program.^43^ Geometry optimization and RESP charge (Merz-Kollman scheme) calculations were performed at the B3LYP/6-31G* level of theory in Gaussian 16.^44–46^ The obtained NCP-NSD2 complex was then solvated in TIP3P water^47^ with NaCl to neutralize the charge and achieve an effective salt concentration of 150 mM. The resulting system consisted of ≈485,000 atoms within a solvation box of size 165 Å *×* 180 Å *×* 180 Å. All the models were prepared using the Leap program in AMBER. ^39^ Similarly, the E1099K, T1150A, and E1099K/T1150A mutants that are reported to hyperactivate the methylation activity of the NSD2 enzyme,^20,26^ were also prepared. Finally, all the systems were subjected to all-atom MD simulations.

### Simulation Details

All the simulations were carried out using the protocol reported in our previous publication.^6,7^ In summary, all the systems including WT and mutated NSD2 systems, comprising the entire nucleosome (DNA, NSD2, and histones), were minimized in two steps: the protein and the nucleosome were strongly restrained (500 kcal mol^−1^ Å^−2^) for the first 5000 steps of minimization, and the latter 10,000 steps of minimization were carried out without any restraint. Upon minimization, the temperature was gradually raised to 300 K for 1.5 ns in the NVT ensemble using a Langevin dynamics with a collision frequency of 1 ps^−1^. During this heating, the protein backbone was restrained at 10 kcal mol^−1^ Å^−2^. The systems were then switched to the NPT ensemble using a Langevin thermostat ^48^ and a Berendsen barostat,^49^ and equilibrated for 4 ns (2 ns with 10 kcal mol^−1^ Å^−2^ restraints and 2 ns without restraints) before production. The simulations were performed with a time step of 2 fs, and snapshots were saved every 50 ps. The cutoff distance of 12 Å was chosen for all the non-bonded interactions, and all bonds involving hydrogen atoms were constrained with the SHAKE algorithm.^50^ The long-range electrostatic interactions were treated using the particle mesh Ewald (PME) method.^51^ Details of the systems and simulation times are shown in **SI Table S1**. MD production simulations were run for 500 ns and were repeated five times with the same initial conformation and different initial velocities.

The simulations were carried out with the GPU-accelerated version of the pmemd engine in the AMBER20 package. ^39^ The obtained trajectories were visualized using VMD ^52^ and the images were generated using ChimeraX.^53^ The root mean square deviation (RMSD) and root mean square fluctuations (RMSF) of backbone atoms was calculated taking the first frame as reference using CPPTRAJ.^54^ Hydrogen bond and contact frequency analyses were performed by taking a frame every 250 ps using VMD.^52^ The hydrogen bond between two polar residues was considered when they met the distance condition of ≤ 4 Å and angle condition of ≤ 35^*◦*^. An in-house TCL script was used to calculate the contact frequency between the desired residue and residues within 4.5 Å.

### Time-lagged Independent Component Analysis and Markov State Modeling

tICA was performed using pairwise distances between C*α* atoms of 30 residues, including the autoinhibitory loop, residues around the mutation site, and H3 binding sites in the NSD2 complex **(Fig. S1)** across all trials. This provided a total of 276 features. The analysis was performed using MDTraj^55^ and PyEmma.^56^ A lag time of 1 ns was chosen using the implied timescale (ITS) test to ensure stable kinetic separation of slow collective modes while preserving sufficient statistical sampling.

The Markov state models (MSMs) were constructed in the collective variable (CV) space obtained from tICA employing PyEMMA version 2.5.12.^56^ A lag time of 0.5 ns was selected based on the plateau of the slowest timescales in the ITS test, indicating approximate Markovianity of the underlying dynamics. To construct the MSMs, we clustered the tICA space into 400 microstates using k-means, which were eventually assigned to macrostates using the robust Perron cluster cluster analysis (PCCA+) algorithm.^57,58^ The mean first-passage times (MFPTs) were used to identify the transition rates between the conformational states. The transition path theory was employed to determine transition pathways and their associated fluxes.^59^ The dominant pathways, states, and total percentages are provided for each system. We also computed free-energy surfaces (FESs) for each case using *F* = −*k*_B_*T* log *P*, where the probability distribution *P* in the CV space was calculated using kernel density estimation.

## Results and Discussion

### Oncogenic Mutations Do Not Alter the Global Dynamics of the Enzyme

Five independent simulations were performed for the NSD2-NCP complex, WT, and three oncogenic mutant systems, totaling 10 *µ*s of simulation time. All five system trajectories were analyzed together to understand overall conformational dynamics and stability. Surprisingly, despite the functional hyperactivity of the mutants, their overall structural dynamics was indistinguishable from those of the WT. The RMSD analysis **(SI Fig. S2)** demonstrates that DNA and histones are the most flexible components. The flexibility of the DNA could be attributed to its extended termini, whereas the flexibility of the histones arises from the inherently dynamic nature of histone tails. ^6,7^ This is evident by comparison of RMSDs of ordered regions of histone (Histone_*ordered*_), which has RMSD ≈1 Å in all the systems throughout the simulations. The RMSD comparison for NSD also shows that the dynamics is similar across all systems, suggesting that the mutants do not alter the enzyme’s global stability. To understand the residue-level impact of oncogenic mutations, we performed RMSF analysis over the trajectories. Unlike RMSD analysis, RMSF shows a slight increase in fluctuations of the residues near the histone binding site (residues 1120–1150) and regulatory loops (residues 1180–1190) upon mutation **(SI Fig. S3)**. However, the differences were not pronounced, and the observed changes were inconsistent across the four systems, suggesting that the mutations do not induce significant destabilization or conformational changes.

### tICA and MSMs Unveil Altered Dynamics and Transition Kinetics

The unperturbed stability of the oncogenic system suggests that the mechanistic basis of hyperactivity lies in subtle alterations to the conformational landscape and dynamic coupling within the enzyme-substrate complex, which are difficult to capture by global fluctuation metrics. To uncover these subtle yet functionally critical changes in conformational dynamics, we employed tICA. Using distance-based features calculated from simulation trajectories (the region from which features were obtained is shown in **SI Fig. S1)**, we projected the first two independent components (ICs, namely *z*_1_ and *z*_2_) and constructed the FES in the CV space for each system (**Fig. 2A**). The FESs reveal notable differences in conformational changes between WT and the mutant variants.

**Figure 2:**
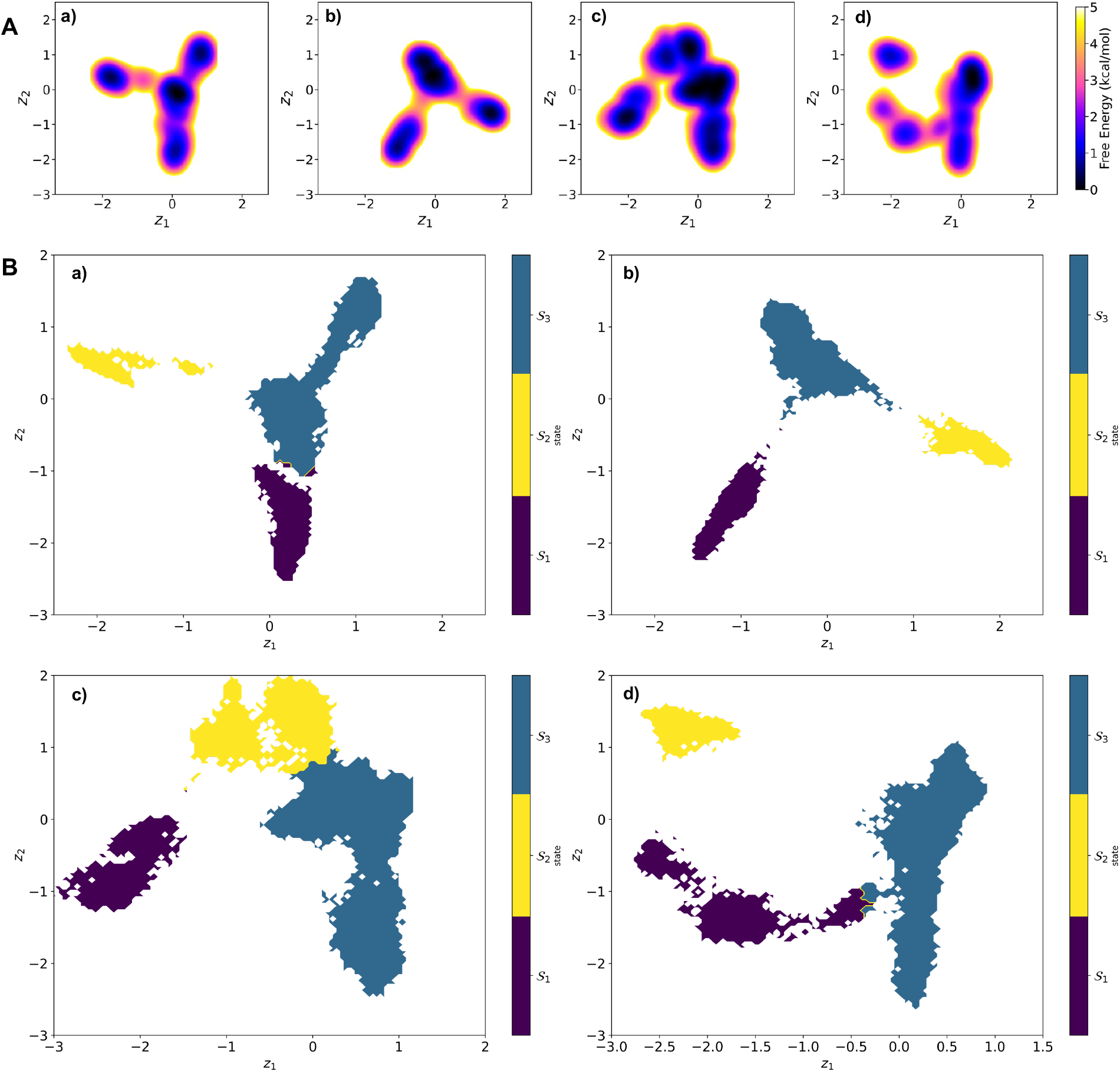
(A) Free-energy surface and (B) macrostate assignments shown on the tICA space of (a) WT, (b) E1099K, (c) T1150A, and (d) E1099K/T1150A.

In the WT system, the FES (**Fig. 2A**, panel (a)) exhibits a Y-shaped landscape, with one basin on the left and multiple low-energy basins extending outwards from the center of the CV space along *z*_1_ axis. These minima are interconnected by moderate energy barriers (<3 kcal/mol) suggest that the interconversion between these states is facile, whereas transitions across *z*_2_ require high-energy barriers, resulting in a conformational ensemble that is diverse yet dynamically restricted. In contrast, the E1099K mutant displays a more localized FES (**Fig. 2A**, panel (b)), with minima positioned at the vertices of the triangle. Compared to WT, this conformational landscape appears to be restrictive and indicative of the stabilization of the specific conformation(s). The T1150A mutant (**Fig. 2A**, panel (c)) exhibits notably heterogeneous ensemble among all the systems, with multiple basins distributed across the CV space. The T1150A mutant has a metastable state at the origin, similar to WT. However, it has broader minima, and transitions along *z*_1_ appear to have a negligible energy barrier. This enhanced delocalization implies that interconversion between the conformational states is energetically favorable and may underlie its modest catalytic enhancement through the proposed allosteric mechanism.^9^ The FES of the double mutant (E1099K/T1150A) (**Fig. 2A**, panel (d)), interestingly exhibits the CV space similar to the WT system, but has transition regions whose FES has been noticeably fragmented compared to WT. In addition, it is also populated in the lower-left corner of the CV space, with higher energy barriers. The coexistence of peripheral minima with altered connectivity suggests that the system is trapped by kinetic bottlenecks, hindering inter-state transitions.

Building upon the tICA projections, we employed MSMs to partition the conformational space into metastable macrostates and investigate the structural and kinetic properties of NSD2 dynamics. The tICA space was discretized using k-means clustering (400 microclusters, **SI Fig. S4**), and macrostates were assigned as shown in **Fig. 2B**. We further constructed the kinetic networks for all systems to summarize the obtained kinetic information, transition conformations, and macrostate probabilities (**Fig. 3A**). These plots illustrate the connectivity and MFPT between macrostates, with distinct transition topologies across the variants. To directly assess the catalytic readiness of each macrostate, we analyzed the distribution of SAM-H3K36 distances within each macrostate across all simulation frames. These distributions are summarized in **Fig. 3B** and provide a direct structural metric to determine the catalytic readiness of the configuration. Distance-based classifications are defined as ≤ 6 Å (near-attack state), 6–8 Å (catalytically competent state), and >8 Å (catalytically dormant state). In the WT system (**Fig. 3B**, panel (a)), macrostates S1 and S2 show broad distributions centered at the longer distances, confirming disengaged configurations with SAM–H3K36 distances >8 Å, while macrostate S3 showed a more compact arrangement (average distance 7.64 *±* 0.82 Å). Despite S3 accounting for ≈ 66% of the population, transitions from S1 or S2 to S3 were slow (MFPT >2.5 *µ*s), whereas transitions out of S3 occurred relatively rapidly (**Fig. 3A**, panel (a)). This kinetic asymmetry suggests that although the WT enzyme occasionally accessed the catalytically competent state (distances 6 to 8 Å), it could not maintain it for extended periods, explaining the predominantly catalytically dormant behavior. The E1099K mutant exhibited markedly altered kinetics with all macrostates having population in the near attack state (distance ≤ 6 Å) (**Fig. 3A** and B, panel (b)). Macrostates S1 and S3, in particular, which accounted for almost 80% of the total population, has ≥50% population in catalytically competent state. Interestingly, the transition to the macrostate S3 was significantly fast (as low as 450 ns). However, consistent with the high energy transition barriers in the FES discussed above, the residence time within this state was very long. This redistribution of flux toward catalytically favorable states, along with localized energy minima in the CV space, aligns with the enhanced methylation activity observed experimentally.

**Figure 3:**
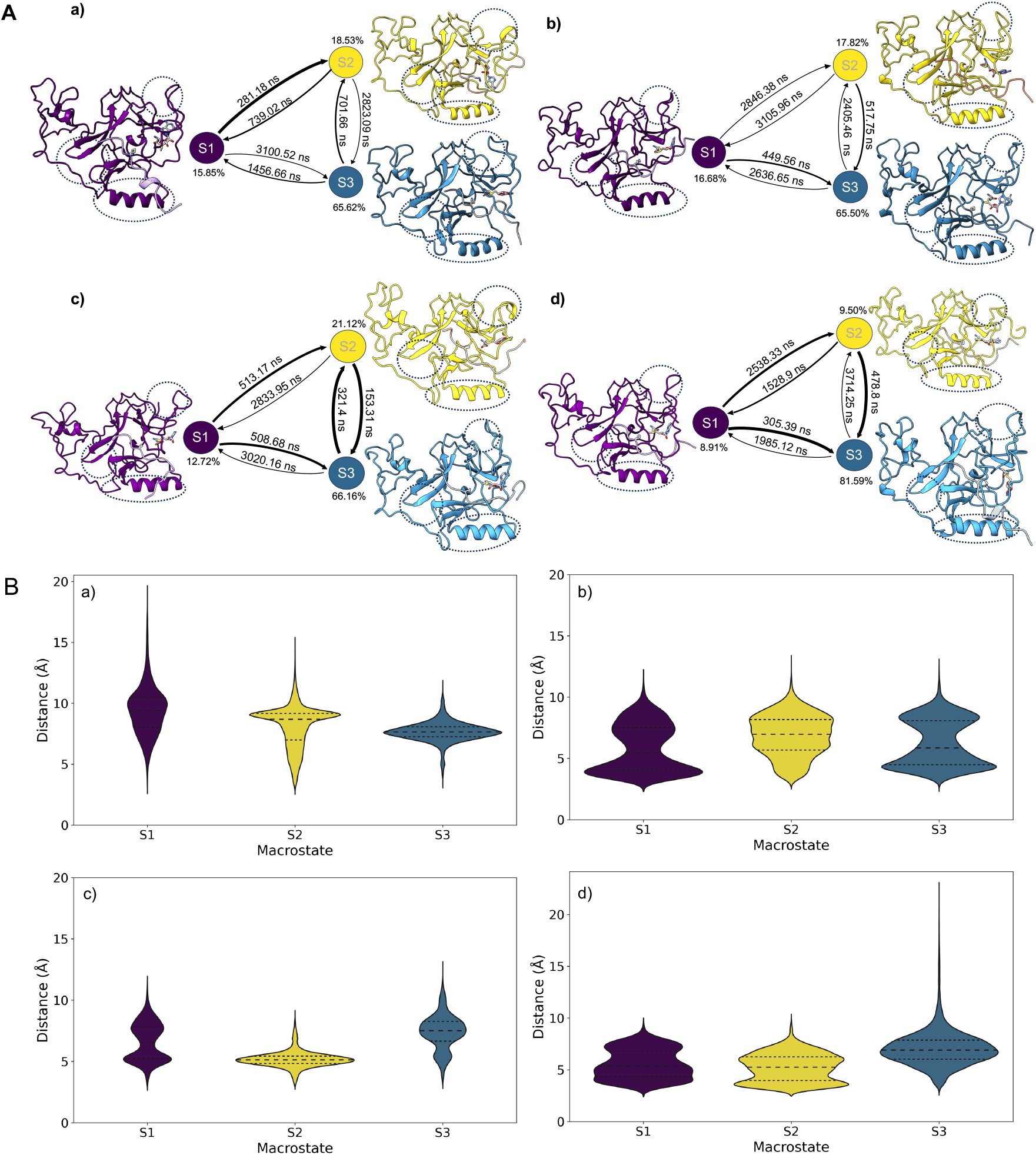
(A) Kinetic networks of conformational states and (B) violin plots of SAM–H3K36 distance distributions across MSM-derived macrostates for (a) WT, (b) E1099K, (c) T1150A, and (d) E1099K/T1150A. The representative conformations extracted from the centers of each macrostate are shown next to each state, along with their population percentages. The thickness of the arrows connecting to macrostates is proportional to the transition rate and is labeled with their respective MFPT values in ns. The median and quartiles are displayed in the violin plots to illustrate the distribution of the data within each macrostate. The distance based definitions of states are: near attack (≤ 6 Å), catalytically competent (6 to 8 Å), and catalytically dormant (>8 Å).

The distributions between macrostates of the T1150A mutant (**Fig. 3B**, panel (c)) were the narrowest. The macrostate S2 almost entirely represented the near-attack conformation, with the other two states also having sizable populations in that conformation. As observed in the FES, this mutant displayed the most dynamic behavior (**Fig. 3A**, panel (c)), with transitions out of the near-attack geometry (macrostate S3 and S1) occurring much faster (as quickly as ≈150 ns). This rapid dynamical exchange among active conformations is consistent with the delocalized FES. The E1099K/T1150A double mutant showed the significant kinetic reorganization (**Fig. 3A**, panel (d)) with macrostate S3 dominating at approximately 82% of the total population. Although this macrostate had a small population (≈0.5%) in the disengaged state, the majority of the conformational ensemble is primed for catalysis. Furthermore, transitions toward S3 were faster than exchanges within the macrostates S1 and S2, indicating that the enzyme efficiently accesses and maintains productive conformations without compromising global stability. This structural view of the tICA and MSMs, along with kinetic networks, supports the interpretation that oncogenic mutations in NSD2 remodel the conformational landscape to promote persistent, catalytically competent substrate engagement, offering a mechanistic basis for their observed hyperactivity.

### Oncogenic Mutations Induce Hyperactivity through Reorganization of Catalytic Interface

Having established through tICA and MSMs that oncogenic mutations systematically remodel the conformational landscape to favor catalytically competent states, we next examined how this manifests in the overall distribution of SAM-H3K36 distances across the entire simulation. It should be noted that we omitted samples with extremely large microstate distances. These outliers represented ≤0.2% of the total data. As shown in **Fig. 4A**, both the WT and single mutant systems exhibited bimodal distance distributions spanning 2.5 Å to 14 Å, indicative of discrete conformational ensembles. The WT system displayed a broad distribution with two distinct peaks at 8 Å and 9.5 Å, suggesting that the enzyme predominantly occupies the catalytically dormant state. In contrast, the single mutants transpired markedly narrower distance distributions confined within ≈2 Å, with almost no population beyond 10 Å. Moreover, the highest probability densities of these systems were found to be within the catalytically active state (<6 Å). This suggests that mutants suppress the distal substrate conformations and confine SAM at the near-attack geometry, thereby promoting enzymatic activity. Surprisingly, the double mutant shows a wide single distribution with the highest probability around 6.5 Å, suggesting reorganization of the enzyme that coerces SAM into the near-attack conformation. To further quantify the enzymatic activity, we computed the percentage of simulation time during which the H3K36-SAM distance remained below 6 Å, a threshold corresponding to the near-attack geometry (the cryo-EM distance is 6.23 Å, **Fig. 4A**, right table). The examination of results implicates the stronger activation effect of E1099K oncogenic mutation than T1150A, which is in accordance with the catalytic activity reported by Sato et al.^20^ Moreover, in a recent report by Khella et al.,^9^ hyperactivity of T1150A mutation was observed only during the LYS dimethylation to trimethylation step through a proposed allosteric effect.^1,9^ In the double mutation, the reduced occupancy of the active state likely arises from competing structural influences of both mutations, where E1099K drives local activation while T1150A introduces conformational rearrangements that partially offset the cooperative alignment of SAM and H3K36, resulting in an intermediate catalytic activity.

**Figure 4:**
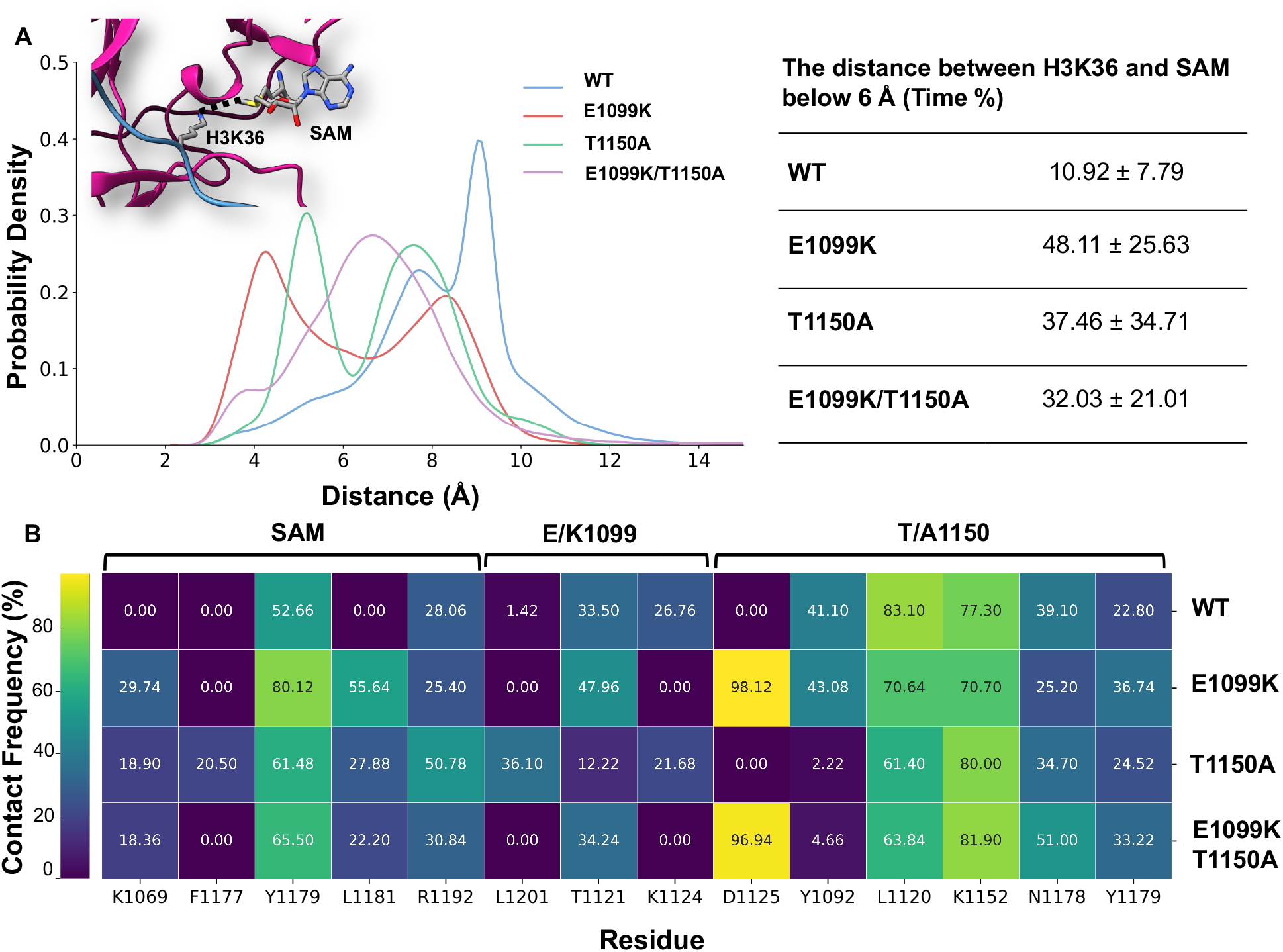
(A) Probability distribution of the distance between *ε*-N of H3K36 and methyl carbon of SAM during the simulations. The probability is plotted for all five trials together. The definition of distance is shown in inset. Right table shows the percentage of simulation time when the SAM-H3K36 distance was below 6 Å. (B) Heat map of SAM, E/K1099, and T/A1150 contact frequency in NSD2 in the % of the simulation time in five trials of WT and mutant systems.

To gain mechanistic insight into the influence of oncogenic mutations on catalytic geometry, we then examined how SAM engages in the active state across four systems. In the catalytically engaged conformation, the donor methyl group of SAM is oriented toward the *ε*-nitrogen of H3K36, to enable methyl transfer (**SI Fig. S5 A and B**). This reactive pose is supported by a tightly choreographed network of interactions of SAM that includes the *π*-stacking of ribose with F1117 and a hydrogen bond with Y1118, while the adenosine ring and carboxylate tail are anchored by K1073, W1075, R1138, M1140, N1141, and Y1179 (**SI Fig. S6**). Residues W1075, Y1118, and N1141 are also conserved in NSD1 and NSD3. Furthermore, Y to F mutations in some methyltransferases have been shown to change the product specificity, ^1,20^ suggesting the functional importance of this SAM-binding architecture in LYS methylation. Despite this conservation, the quantitative analysis revealed systematic differences in SAM coordination. The hydrogen bonding of SAM with W1075 and N1141 was increased in mutant systems, suggesting enhanced stabilization of SAM in its catalytically engaged orientation more effectively than WT (**SI Table S2**). Contact frequency analysis further revealed that mutant systems establish additional stabilizing interactions involving Y1179, L1181, and K1069 (**Fig. 4B**). The emergence of K1069 contact is significant given its proximity to both the SAM carboxylate and W1075, which also form a persistent hydrogen bond with SAM. The Y1179 has also been implicated in proper alignment of the substrate within the catalytic pocket, and the strengthened contact in the mutants likely reinforces this positioning, favoring the methyl-transfer geometry.^60,61^ Together, these residues appear to form a cooperative electrostatic network that stabilizes the cofactor in optimal geometry for efficient methyl transfer. The T1150A mutant, however, displayed a uniquely remodeled interaction pattern, where SAM formed additional contacts with F1177, R1192, and L1201 near the post-SET region. These interactions, along with faster dynamical exchange among active conformations, could further reflect an allosteric pathway that reorganizes loop dynamics to stabilize the active conformation, consistent with the moderate active conformation observed in the distance analysis for this mutant.

Examination of the contact profile around the mutation site further underscored how local interactions propagate toward the catalytic center. In the E1099K mutant, the charge reversal perturbed the local electrostatic environment, which resulted in a modest increment in hydrogen bonding with *α*-helix. However, the contact frequency analysis revealed that the substitution abolished contact with K1124 but introduced persistent interaction with D1125 and increased contact with T1121 (**Fig. 4B**, **SI Fig. S7 A**). Given the close proximity of these residues to the SAM binding site (F1117 and Y1118) and the histone substrate, this electrostatic reorganization likely plays a key role in promoting optimal alignment of both SAM and histone within the active site. This was further observed in the distance analysis, where E1099K showed a narrow distribution and was in the active conformation for more than 48% of the simulation time. The introduction of T1150A mutation produced markedly different changes, characterized by a decrease in the contact of residues Y1092 and L1120 (**Fig. 4 B**, **SI Fig. S7 B**). This is particularly notable in the contact with Y1092 in the T1150A and E1099K/T1150A systems, where it was reduced to <5%. This observation again corroborates the findings of Khella et al.,^9^ who reported a similar reduction in L1120-Y1092 contacts and showed that this contact emancipation increases the volume of the catalytic site and enhances activity. Overall, these contact rearrangements delineate how oncogenic mutations remodel the local interaction network to enhance cofactor stabilization and substrate positioning, thereby pre-organizing the catalytic ensemble for increased methyl-transfer efficiency.

In summary, these observations demonstrate that while the SAM-binding core remains structurally conserved across wild-type and mutant NSD2, oncogenic mutations systematically remodel the surrounding interaction network to enhance catalytic efficiency through complementary mechanisms. The E1099K mutation establishes new electrostatic contacts (D1125, T1121) that propagate stabilizing effects to the SAM-binding pocket, while T1150A relieves steric constraints (loss of Y1092, L1120 contacts) to expand the catalytic volume and enable the allosteric flexibility observed in tICA and MSMs. Both mutations converge on a common outcome, strengthened SAM stabilization through enhanced interactions with the W1075-K1069 cooperative network, coupled with reorganization of the aromatic cage (Y1092, Y1179) that optimizes substrate positioning. The synergistic effect in the E1099K/T1150A double mutant reflects the integration of both mechanisms: electrostatic pre-organization of the active site combined with enhanced conformational sampling, resulting in the hyperactive yet dynamically balanced enzyme observed experimentally. These molecular-level insights, when integrated with the conformational landscape remodeling revealed by tICA and MSMs, provide a comprehensive mechanistic framework explaining how distant oncogenic mutations can profoundly enhance NSD2 catalytic activity, not through alteration of fundamental cofactor binding modes, but through subtle yet systematic optimization of the catalytic geometry and kinetic accessibility of productive states.

## Conclusion

In this work, we investigated the structural and dynamic effects of oncogenic mutations in NSD2 bound to the nucleosome, focusing on the WT, E1099K, T1150A, and E1099K/T1150A systems. The tICA and MSMs revealed striking alterations in the energetic and kinetic landscapes of the mutant systems, favoring prolonged, more frequently sampled catalytically competent states and reduced SAM-LYS distances. The mutations systematically rewired the network of intramolecular contacts around the catalytic site, histone-binding interface, and SAM-binding pocket. This contact reorganization enabled more efficient substrate engagement and proper alignment of catalytic residues, establishing the structural foundation for methyltransferase hyperactivity. Collectively, our results demonstrate that oncogenic NSD2 mutations exert their hyperactive effects not through subtle yet profound rewiring of the enzyme’s internal dynamics. The mutations achieve a balance between local reorganizations at key functional sites and global conformational flexibility, allowing the enzyme to maintain catalytic readiness while preserving the global stability necessary for substrate recognition and turnover. These findings provide crucial molecular insights into how NSD2 dysregulation drives oncogenesis and may inform the rational design of allosteric inhibitors targeting the enzyme’s conformational dynamics.

## Data Availability

The input parameter and coordinate files for MD simulations, tICA features, and pipelines to build tICA and MSMs are available at Zenodo (DOI: https://doi.org/10.5281/zenodo.20422910).

## Author Contributions

H.T. designed the project; H.T. and J.R. supervised the project; T.S. performed the simulations and analyses; S.H. collected simulation data and analysis; All authors reviewed data and wrote the manuscript together.

## Acknowledgement

This work was supported by National Institutes of Health grant R35 (GM155106) and a startup grant from the University of Texas at Dallas to H.T. The content is solely the responsibility of the authors and does not necessarily represent the official views of the National Institutes of Health. Authors also acknowledge the Texas Advanced Computing Center (TACC) at the University of Texas at Austin and the Office of Information Technology and Cyber Infrastructure Research Computing (CIRC) department and the High Performance Computing at the University of Texas at Dallas (HPC@UTD) for their technical assistance and providing the computing resources that have contributed to this research. T.S acknowledges University of Texas at Dallas office of Research for Graduate Research and Cancer Education (GRACE) Fellowship and the Polish National Agency for Academic Exchange (NAWA) and Nicolaus Copernicus University for the PROM fellowship. J.R. acknowledges funding from the National Science Center in Poland (Sonata 2021/43/D/ST4/00920, “Statistical Learning of Slow Collective Variables from Atomistic Simulations”).

## Supporting Information Available

The following information is available free of charge.

- Tables: simulation times and average H-bond occupancy between residue pairs.
- Figures: features selected for tICA and MSMs; time evolution of RMSD; Δ RMSF of backbone residues in NSD2 enzyme; k-means clustering in the tICA space; representative conformations of SAM bound in NSD2; residues interacting with SAM; representative residues interacting with K1099 or T1150.

