## Supplementary Information for "Decoding the Conformational Dynamics and Hyperactivity of Histone H3K36 N-Methyltransferase in Oncogenic Mutations *via* tICA and Markov State Modeling"

Table S1: Simulation Times

| Simulation time/trial (ns) | | | | Total Simulation time |
|---|---|---|---|---|
| WT | E1099K | T1150A | E1099K | |
| 500 × 5 | 500 × 5 | 500 × 5 | 500 × 5 | 10 $\mu$s |

Table S2: Average hydrogen-bond occupancy (%) between residue pairs, representing the fraction of simulation time in which a hydrogen bond was present. Values are averaged over five independent trials.

| Residue | Residue | WT | E1099K | T1150A | E1099K/T1150A |
|---|---|---|---|---|---|
| SAM | W1075 | 41.16 ± 24.44 | 62.37 ± 28.23 | 53.71 ± 33.54 | 44.39 ± 18.33 |
| | Y1118 | 45.25 ± 21.18 | 42.73 ± 36.23 | 40.15 ± 22.53 | 41.54 ± 16.24 |
| | N1141 | 32.54 ± 30.73 | 43.15 ± 36.23 | 56.95 ± 31.22 | 63.73 ± 34.30 |

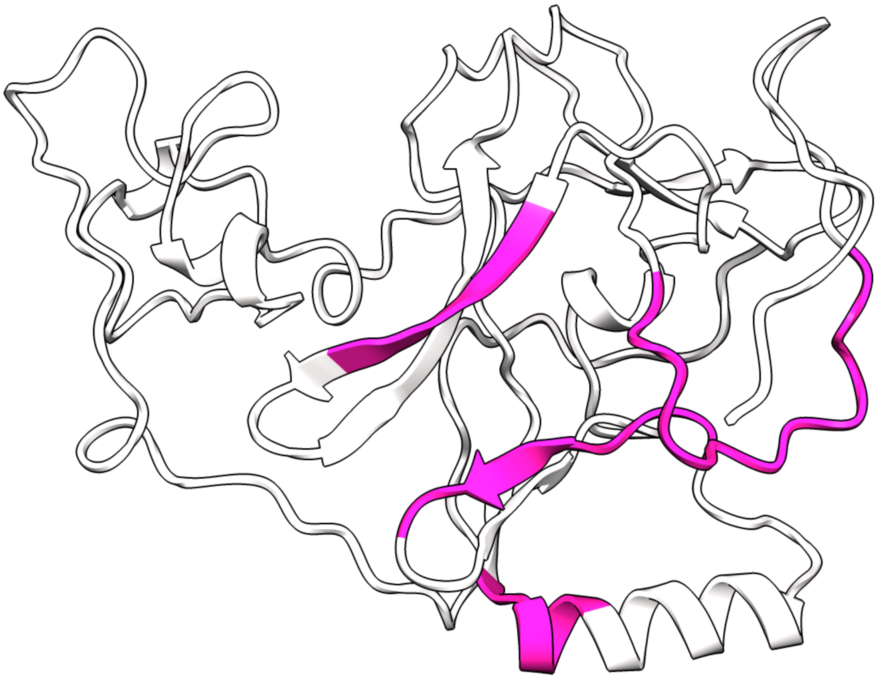

Figure S1: The region selected to obtain pairwise distance to perform tICA and MSM analysis to study slow dynamics. The pairwise distances of C$\alpha$ carbons of residues colored magenta were used as a feature of the analysis.

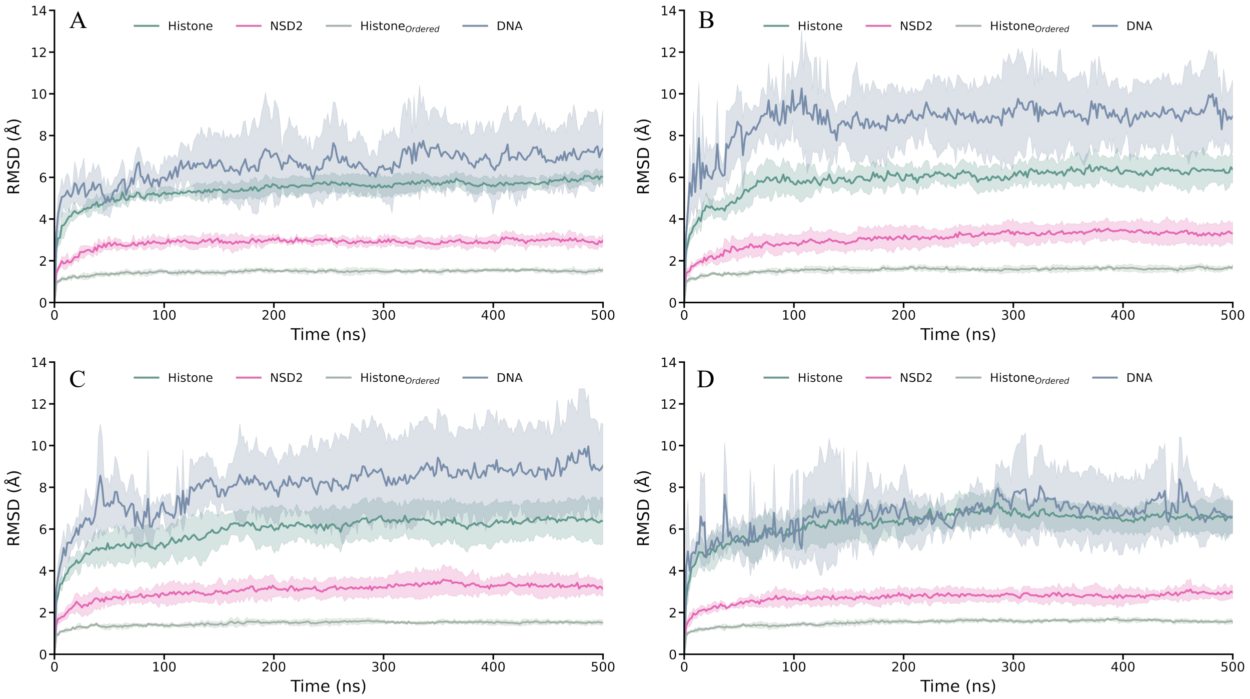


Figure S2: Time evolution of RMSD of (A) WT, (B) E1099K, (C) T1150A and (D) E1099k/T1150A systems. The solid line represents the average of five trials and shaded regions shows standard deviation. Color codes: Green - Histone, Magenta - NSD2, Olive - Ordered region of Histones, and Navy - DNA.

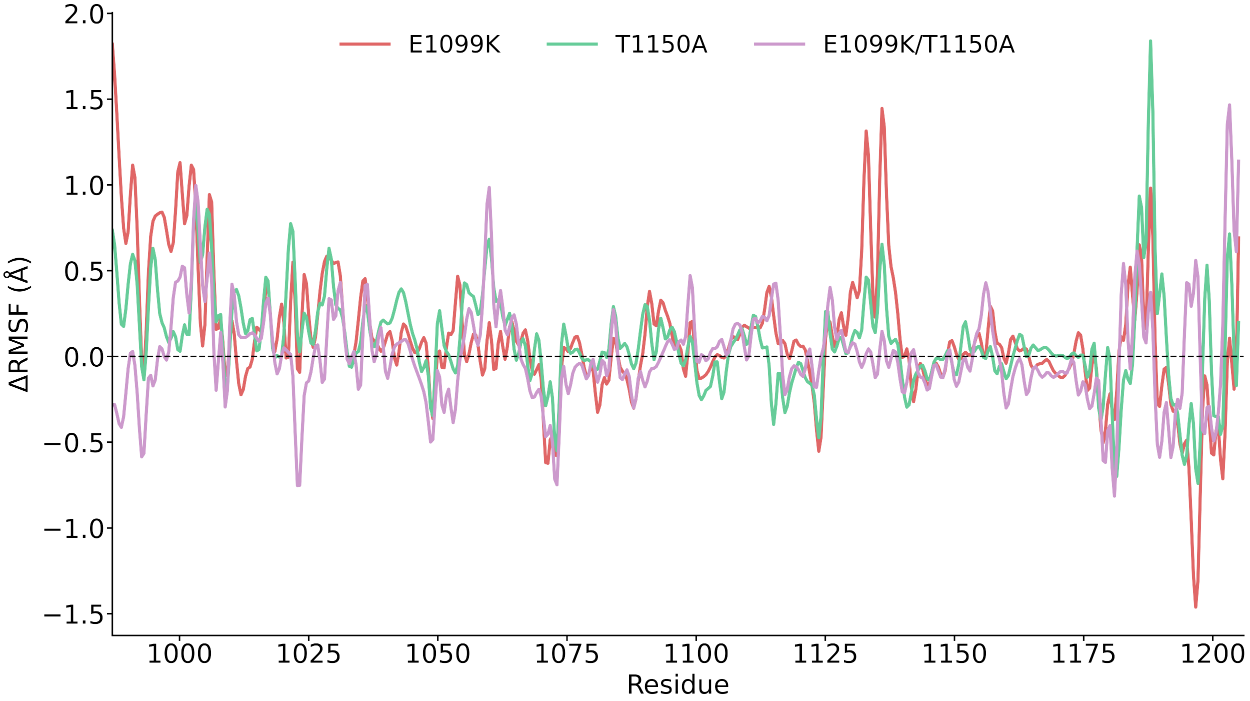


Figure S3: ΔRMSF of NSD2 complex in E1099K (red), T1150A (green), and E1099K/T1150A (magenta) systems. The plotted line represents the average RMSF of five trials. ΔRMSF = $\text{RMSF}_{mutant}$ - $\text{RMSF}_{WT}$.

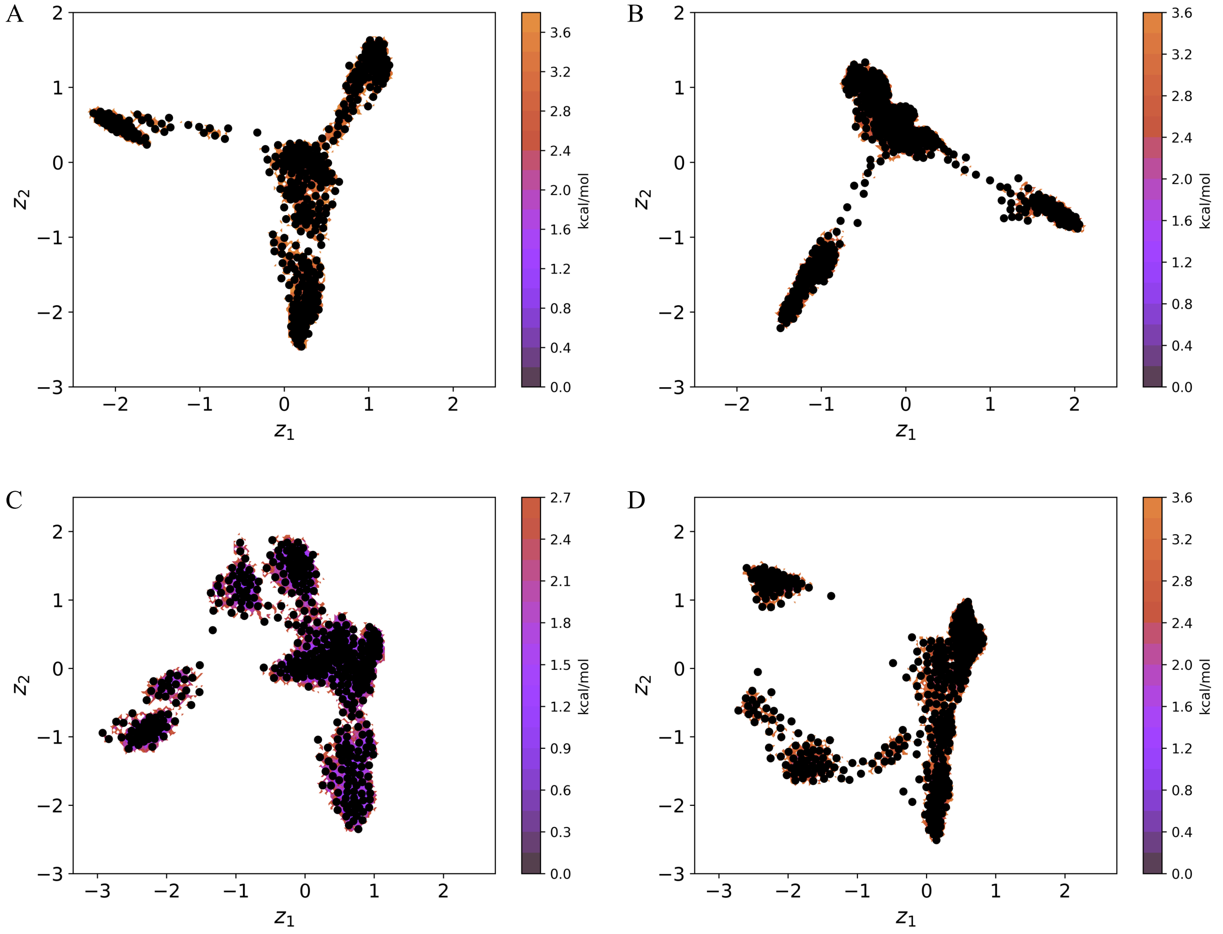


Figure S4: The k-means cluster centers overlaid on the tICA free energy surfaces for (A) WT, (B) E1099K, C) T1150A, and D) E1099K/T1150A systems. The plots show the distribution of conformations along the top two time-lagged independent components, with cluster centers (black dots) marking the partitioning used for MSM construction. This clustering defines the basis for assigning kinetically meaningful states in the subsequent analysis.

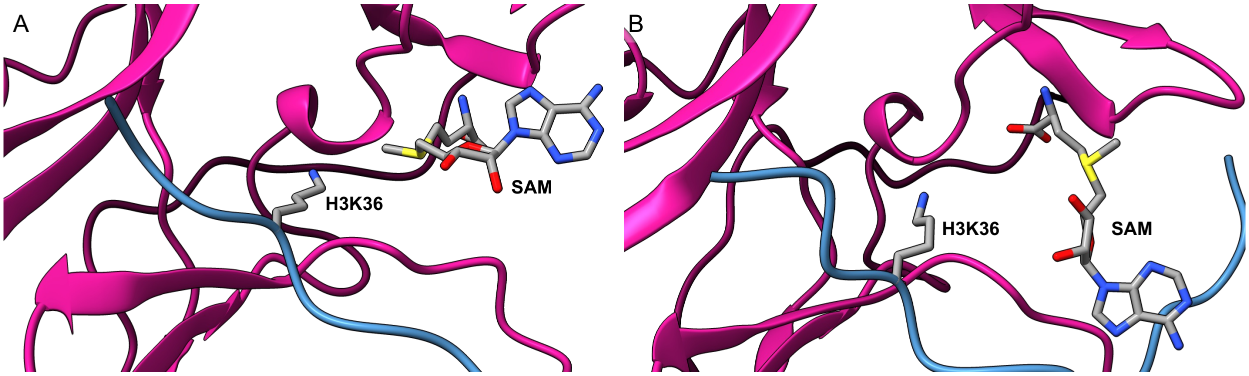


Figure S5: Representative conformations of SAM bound to NSD2: (A) engaged form and (B) disengaged form.

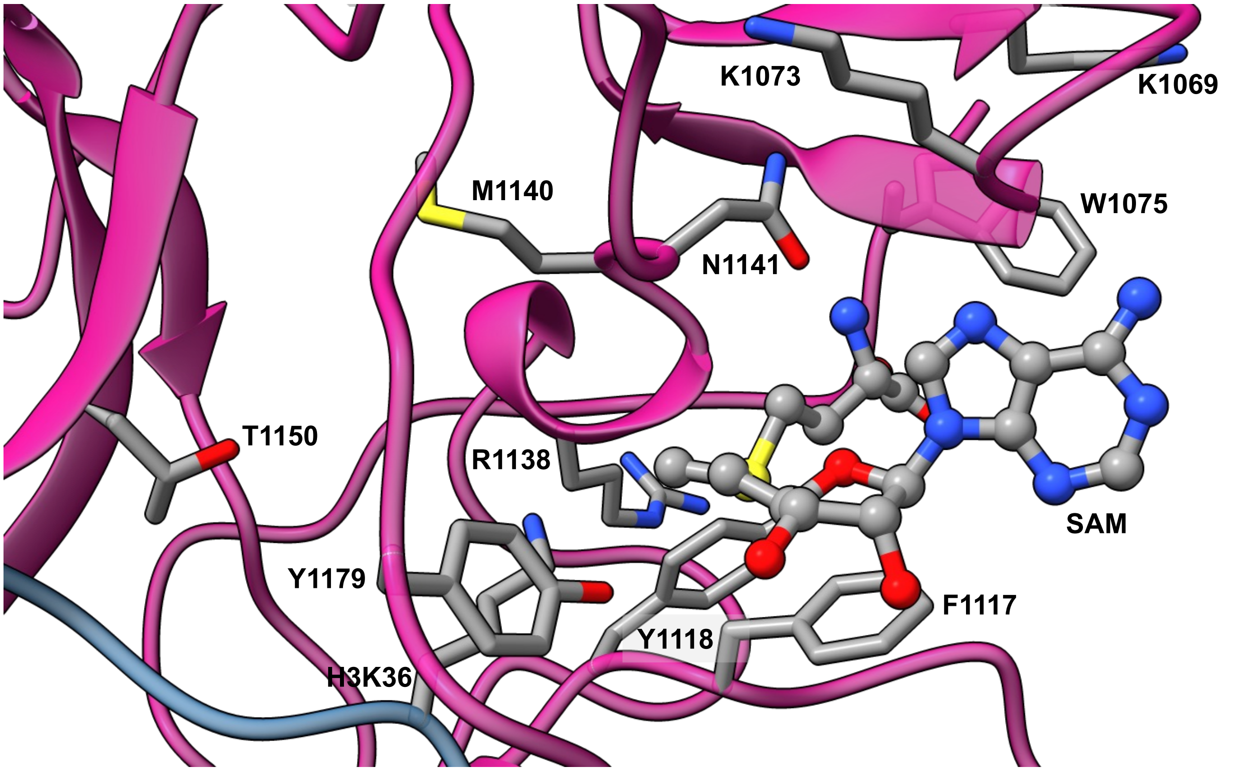


Figure S6: Key residues forming hydrogen bond interactions with SAM in the binding pocket.

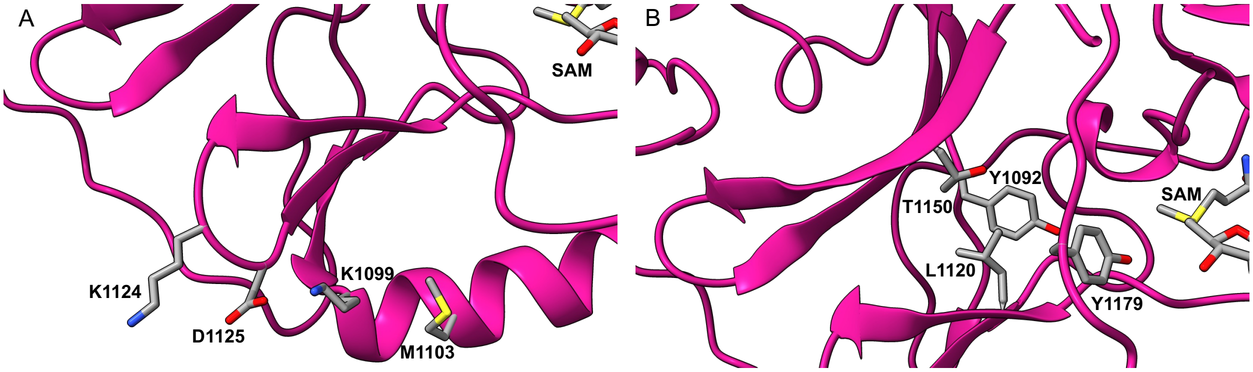


Figure S7: Representative figure to show the residues interacting with (A) K1099 and (B) T1150 in the NSD2 enzyme. Figure was taken from E1099K/T1150A system.
